# Characterization of an α-glucosidase enzyme conserved in *Gardnerella* spp. isolated from the human vaginal microbiome

**DOI:** 10.1101/2020.05.11.086124

**Authors:** Pashupati Bhandari, Jeffrey P. Tingley, David R. J. Palmer, D. Wade Abbott, Janet E. Hill

## Abstract

*Gardnerella* spp. in the vaginal microbiome are associated with bacterial vaginosis, a dysbiosis in which a lactobacilli dominant microbial community is replaced with mixed aerobic and anaerobic bacteria including *Gardnerella* species. The co-occurrence of multiple *Gardnerella* species in the vaginal environment is common, but different species are dominant in different women. Competition for nutrients, particularly glycogen present in the vaginal environment, could play an important role in determining the microbial community structure. Digestion of glycogen into products that can be taken up and further processed by bacteria requires the combined activities of several enzymes collectively known as amylases, which belong to glycoside hydrolase family 13 (GH13) within the CAZy classification system. GH13 is a large and diverse family of proteins, making prediction of their activities challenging. SACCHARIS annotation of the GH13 family in *Gardnerella* resulted in identification of protein domains belonging to eight subfamilies. Phylogenetic analysis of predicted amylase sequences from 26 *Gardnerella* genomes demonstrated that a putative α-glucosidase-encoding sequence, CG400_06090, was conserved in all species in the genus. The predicted α-glucosidase enzyme was expressed, purified and functionally characterized. The enzyme was active on a variety of maltooligosaccharides over a broad pH range (4.0 - 8) with maximum activity at pH 7. The *K_m_*, *k*_cat_ and *k_cat_/K_m_* values for the substrate 4-nitrophenyl α-D-glucopyranoside were 8.3 μM, 0.96 min^-1^ and 0.11 μM^-1^min^-1^ respectively. Glucose was released from maltose, maltotriose, maltotetraose and maltopentaose, but no products were detected on thin layer chromatography when the enzyme was incubated with glycogen. Our findings show that *Gardnerella* spp. produce an α-glucosidase enzyme that may contribute to the complex and multistep process of glycogen metabolism by releasing glucose from maltooligosaccharides.

## Introduction

*Gardnerella* spp. in the vaginal microbiome are hallmarks of bacterial vaginosis, a condition characterized by replacement of the lactobacilli dominant microbial community with mixed aerobic and anaerobic bacteria including *Gardnerella*. This dysbiosis is associated with increased vaginal pH, malodorous discharge and the presence of biofilm (1). In addition to troubling symptoms, the presence of abnormal vaginal microbiota is associated with increased risk of HIV transmission and infection with other sexually transmitted pathogens such as *Neisseria gonorrhoeae* and *Trichomonas* spp. (2, 3). Historically, *Gardnerella* has been considered a single species genus. Jayaprakash *et al*. used cpn60 barcode sequences to divide *Gardnerella* spp. into four subgroups (A-D) (4), and this framework was supported by whole genome sequence comparison (5, 6). More recently, Vaneechoutte *et al*. emended the classification of *Gardnerella* based on whole genome sequence comparison, biochemical properties and matrix-assisted laser desorption ionization time-of-flight mass spectrometry and proposed the addition of three novel species: *Gardnerella leopoldii, Gardnerella piotii* and *Gardnerella swidsinskii* (7).

Colonization with multiple *Gardnerella* species is common, and different species are dominant in different women (8). Understanding factors that contribute to differential abundance is important since the species may differ in virulence (6) and they are variably associated with clinical signs (8, 9). Several factors, including inter-specific competition, biofilm formation and resistance to antimicrobials could contribute to the differential abundance of the different species. Khan *et al*. showed that resource-based scramble competition is frequent among *Gardnerella* subgroups (10). Glycogen is a significant nutrient for vaginal microbiota, but previous reports on the growth of *Gardnerella* spp. on glycogen containing medium are inconsistent (11). Species may differ in their ability to digest glycogen and utilize the breakdown products, which may in turn contribute to determining microbial community structure.

Glycogen is an energy storage molecule, which consists of linear chains of approximately 13 glucose molecules covalently linked with α-1,4 glycosidic linkages, with branches attached through α-1,6 glycosidic bonds (12, 13). A single glycogen molecule consists of approximately 55,000 glucose residues with a molecular mass of ~10^7^ kDa (14). The size of glycogen particles can vary with source; from 10-44 nm in human skeletal muscle to approximately 110-290 nm in human liver (15). Glycogen is deposited into the vaginal lumen by epithelial cells under the influence of estrogen (16), and the concentration of cell-free glycogen in vaginal fluid varies greatly, ranging from 0.1 - 32 μg/μL (17). This long polymeric molecule must be digested into smaller components that can be taken up by bacterial cells and metabolized further (18, 19).

Glycogen digestion is accomplished by the coordinated action of enzymes collectively described as amylases (20). A large majority of these enzymes belong to family 13 within the glycosyl hydrolase class (GH13) of carbohydrate active enzymes (CAZymes) (21). GH13 enzymes are further classified into more than 43 subfamilies based on structure and activity (22). For example, subfamily 23 is primarily composed of α-glucosidases, which are exo-acting enzymes that act on α-1,4 glycosidic bonds from the non-reducing end to release glucose; subfamily 32 is composed of α-amylases, endo-acting enzymes that cleave α-1,4 glycosidic bonds to produce maltose and α-limit dextrin; and subfamily 14 contains debranching enzymes, such as pullulanase, which target α-1,6 glycosidic bonds (15, 23). Limited information is available on glycogen degradation mechanisms in the vaginal microbiome. Although it has been demonstrated that vaginal secretions exhibit amylase activity that can degrade glycogen (19), the role of bacterial amylase enzymes in this process is unknown and there is limited information about glycogen metabolism by clinically important bacteria such as *Gardnerella* spp..

The objective of the current study was to annotate GH13 enzymes in *Gardnerella* spp. and to characterize the activity of a putative α-glucosidase conserved among all *Gardnerella* spp..

## Methods

### *Gardnerella* isolates

Isolates of *Gardnerella* used in this study were from a previously described culture collection (6) and included representatives of *cpn60* subgroup A (*G. leopoldii* (n = 4), *G. swidsinskii* (n = 6), other (n = 1)), subgroup B (*G. piotii* (n = 5), Genome sp. 3 (n = 2)), subgroup C (*G. vaginalis* (n = 6)), and subgroup D (Genome sp. 8 (n = 1), Genome sp. 9 (n = 1)). Whole genome sequences of these isolates had been determined previously (BioProject Accession PRJNA394757) and annotated by the NCBI Prokaryotic Genome Annotation Pipeline (PGAP) (24).

### Identification and functional annotation of putative amylase sequences

Putative amylase sequences were identified in proteomes of 26 *Gardnerella* isolates used in this study based on the PGAP annotations. Multiple sequence alignments of predicted amylase sequences were performed using CLUSTALw with results viewed and edited in AliView (version 1.18) (25) prior to phylogenetic tree building using PHYLIP (26). SignalP v 5.0 (27) and SecretomeP 2.0 (28) were used to identify signal peptides.

Putative amylolytic enzymes in *G. vaginalis* ATCC 14018 and *G. swidsinskii* GV37 (genomes available in the CAZy database) were identified using dbCAN, and predicted sequences were run through the SACCHARIS pipeline. SACCHARIS combines user sequences with CAZy derived sequences and trims sequences to the catalytic domain using dbCAN2 (29). Domain sequences were aligned with MUSCLE (30), and a best-fit model was generated with ProtTest (31). Final trees were generated with FastTree (32) and visualized with iTOL (33). Alignment of CG400_06090 from *G. leopoldii* NR017 to orthologues from *Halomonas* sp. (BAL49684.1), *Xanthomonas campestris* (BAC87873.1), and *Culex quinquefasciatus* (ASO96882.1) was done using CLUSTALw and visualized in ESPript (34). A predicted structure model of CG400_06090 was aligned with *Halomonas* sp. HaG (BAL49684.1, PDB accession 3WY1) (35) to identify putative catalytic residues using PHYRE2 (36). BLASTp (37) alignment of CAZy derived sequences from reference genomes to a database of the predicted proteomes of 26 study isolates was conducted to identify orthologs.

### Expression and purification of the CG400_06090 gene product

Genomic DNA from *G. leopoldii* NR017 was extracted using a modified salting out procedure, and the CG400_06090 open reading frame (locus tag CG400_06090 in GenBank accession NNRZ01000007, protein accession RFT33048) was PCR amplified with primers JH0729-F (5’AT<u>G CAT GC</u>G CAT TAT ACG ATC ATG CTC-3’) and JH0730-R (5’AT<u>G GTA CC</u>T TAC ATT CCA AAC ACT GCA-3’). Underlined sequences indicate *SphI* and *KpnI* restriction enzyme sites. PCR reaction contained 1 × PCR reaction buffer (0.2 M Tris-HCl pH 8.4, 0.5 M KCl), 200 μM each dNTPs, 2.5 mM MgCl_2_, 400 nM each primer, 1 U/reaction of Platinum Taq DNA polymerase high-fidelity (5 units/μL in 50% glycerol, 20 mM Tris-HCl, 40 mM NaCl, 0.1 mM EDTA, and stabilizers) (Life Technologies) and 2 μL of template DNA, in a final volume of 50 μL. PCR was performed with following parameters: initial denaturation at 94 °C for 3 minutes followed by (denaturation at 94 °C, 15 seconds; annealing at 60 °C, 15 seconds and extension at 72 °C, 2 minutes; 35 cycles), and final extension at 72 °C for 5 minutes. Purified PCR products were digested with *KpnI* and *SphI* and ligated into expression vector pQE-80L (Qiagen, Mississauga, ON) digested with same restriction endonucleases. The resulting recombinant plasmid was used to transform One Shot TOP10 chemically competent *E. coli* cells (Invitrogen, Carlsbad, California). Colony PCR was performed to identify transformants containing vector with insert. Insertion of the putative amylase gene in-frame with the N-terminal 6×Histidine tag was confirmed by sequencing of the purified plasmid.

*E. coli* cells containing the plasmid were grown overnight at 37 °C in LB medium with 100 μg/ml ampicillin. Overnight culture was diluted 1:60 in fresh medium to a final volume of 50 mL, and the culture was incubated at 37 °C with shaking at 225 rpm until it reached an OD_600_ of 0.6. At this point, expression was induced with 0.1 mM IPTG and a further 4 hours of incubation was done at 37°C at 225 rpm. Cells were harvested by centrifugation at 10,000 × *g* for 30 mins and the pellet was resuspended in lysis buffer (50 mM NaH_2_PO_4_, 300 mM NaCl, 10 mM imidazole, pH 8.0). Lysozyme was added to 1 mg/mL and cells were lysed by sonication (8 mins total run time with 15 sec on and 30 sec off). The lysate was clarified by centrifugation at 10,000 × *g* for 30 mins at 4 °C and the supernatant was applied to an Ni-NTA affinity column according to the manufacturer’s instructions (Qiagen, Germany). Bound proteins were washed twice with wash buffer (50 mM NaH_2_PO_4_, 300 mM NaCl, 20 mM imidazole, pH 8.0) and eluted with elution buffer (50 mM NaH_2_PO_4_, 300 mM NaCl, 250 mM imidazole, pH 8.0). Eluted protein was buffer exchanged with the same elution buffer but without imidazole using 30 kDa MWCO protein concentrator (ThermoFisher Scientific). Final protein concentration was measured by spectrophotometry.

### Enzyme assay and kinetics

Purified protein was tested for α-glucosidase activity by measuring the release of 4-nitrophenol from a chromogenic substrate 4-nitrophenyl α-D-glucopyranoside as described elsewhere (38). Briefly, 200 μL of enzyme solution (50 nM) was added to 200 μL of 20 mM 4-nitrophenyl α-D-glucopyranoside substrate and the reaction mixture was incubated at 37 °C. A 50-μL aliquot of the reaction mixture was removed and added to 100 μL of 1 M Na_2_CO_3_ solution every 90 seconds up to 7.5 mins (total 6 different time points). The absorbance of this solution was measured at 420 nm, and the amount of 4-nitrophenol released was calculated from a standard curve. To determine activity at different pH values, the substrate was prepared in 50 mM sodium phosphate buffer (pH 4.0-8.0). Kinetic constants were determined for 4-nitrophenyl α-D-glucopyranoside by measuring rate of reaction of enzyme (50 nM) with substrate concentrations of 0.05, 1, 2, 5, 10, 20 and 25 mM at pH 7. Values were fit to the Michaelis-Menten equation, *ν* = *k_cat_* [E]_0_[S]/(*K_m_* + [S]), where *ν* is the observed rate of reaction, [E]_0_ is the initial enzyme concentration, [S] is the substrate concentration, *K_m_* is the Michaelis constant, and *k*_cat_ is the turnover number. Kinetics calculations were performed in GraphPad Prism 8.

### Enzymatic activity on different substrates

The activity of the purified enzyme was examined with different substrates. Briefly, 750 μL of enzyme solution (50 nM) was added to 750 μL of 3 mM maltose, maltotriose, maltotetraose, maltopentaose, or 0.1% (w/v) of maltodextrins (MD 4-7, MD 13-17 and MD 16.5-19.5) or bovine liver glycogen in 50 mM sodium phosphate buffer, pH 6.0, and the reaction mixture was incubated aerobically at 37 °C. Aliquots of the reaction mixture (250 μL) were removed at various time intervals of incubation and were heat-inactivated at 93 °C for 1 minute. The reaction mixtures were centrifuged at 10,000 × *g* for 1 min and supernatant was stored at −20 °C and analyzed by thin layer chromatography (TLC) and high performance anion exchange chromatography (HPAEC) with pulsed amperometric detection (PAD). TLC was performed on silica plates in 1-butanol: acetic acid: distilled water (2:1:1 v/v/v) mobile phase and stained using ethanol: sulfuric acid (70:3 v/v) solution containing 1% (w/v) orcinol monohydrate (Sigma). HPAEC-PAD samples were diluted to 50 μM and resolved on a Dionex PA20 column in a mobile phase of 30 mM NaOH and gradient of 10 mM – 120 mM sodium acetate over 40 min at a flow rate of 0.5 mL min^-1^. Data was analyzed using Chromeleon v6.80 chromatography data system software. Maltooligosaccharide standards ranging from maltose to maltopentaose (Carbosynth) were used for both TLC and HPAEC-PAD analysis; whereas, isomaltotriose, D-panose, and 6-o-a-D-glucosyl-maltose (Carbosynth) were used to identify minor peaks resolved by HPAEC-PAD.

## Results

### Identification of putative amylase sequences

A total of 60 proteins from the 26 study isolates were annotated by PGAP as amylases. After removal of severely truncated sequences, a multiple sequence alignment of the remaining 42 sequences was trimmed to the uniform length of 290 amino acids. Phylogenetic analysis of these predicted amylase sequences revealed five clusters, but only one of these clusters contained orthologous sequences from all 26 isolates (Figure 1). Sequence identities within this cluster were all 91-100% identical. This conserved sequence (corresponding to GenBank accession WP_004104790) was annotated as an GH13 α-amylase (EC 3.2.1.1) in all genomes, suggesting it was an endo-acting enzyme that releases maltooligosaccharides from glycogen.

**Figure 1.**
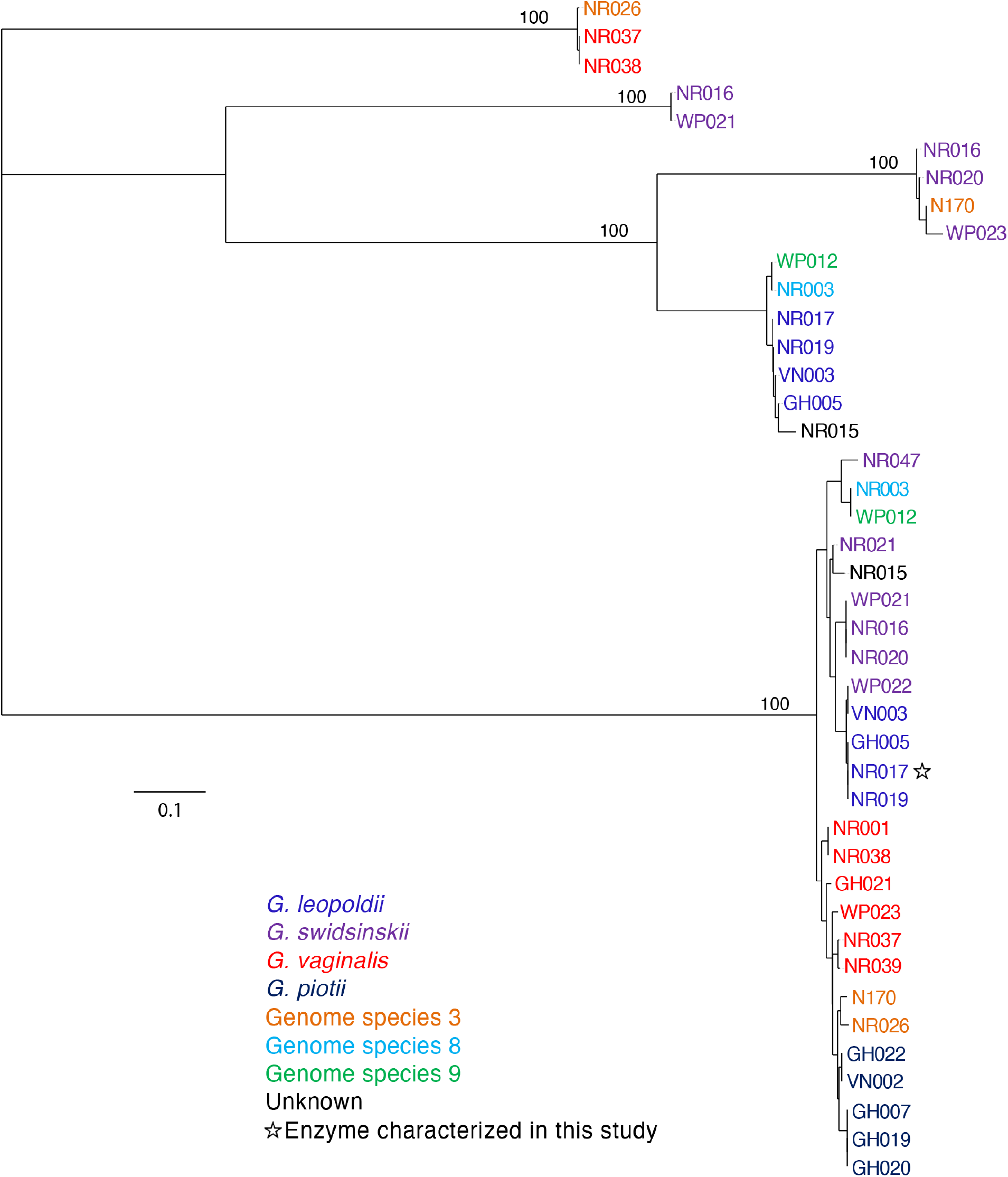
Phylogenetic analysis of predicted extracellular amylases protein sequences from 26 *Gardnerella* genomes. Trees are consensus trees of 100 bootstrap iterations. Bootstrap values are shown for major branch points. Species is indicated by label colour according to the legend.

The GH13 family in the CAZy database is a large, polyspecific family targeting α-glucosyl linkages. The functional diversity of members in this family can make annotation of new sequences challenging. In order to predict the activity of the conserved protein identified among the study isolates, we employed SACCHARIS, which combines CAZyme family trees generated from biochemically characterized proteins with related sequences of unknown function. Currently, two *Gardnerella* reference genomes are available in CAZy: *G. vaginalis* ATCC 14018 and *G. swidsinkii* GV37. GH13 family proteins encoded in these *Gardnerella* genomes were identified and a family tree including all characterized GH13 sequences in the CAZy database was generated (Figure S1). SACCHARIS annotation of the GH13 family in *Gardnerella* resulted in identification of protein domains belonging to eight subfamilies (2, 9, 11, 14, 20, 23, 30, 31 and 32). Alignment with characterized GH13 enzymes from the CAZy database (22) suggested that the conserved amylase identified in the study isolates was an α-glucosidase (EC 3.2.1.20), closely related to subfamilies 23 and 30 of GH13, and not an α-amylase as annotated in GenBank. Subsequent examination of the α-glucosidase with SignalP and SecretomeP indicated that no signal peptide was present, and that the protein was not likely secreted through a Sec-independent pathway.

### Relationship to other α-glucosidases

A representative sequence of the predicted α-glucosidase was selected from *G. leopoldii* (NR017). This sequence (CG400_06090) was combined with functionally characterized members of the GH13 CAZy database using SACCHARIS (Figure 2A). CG400_06090 partitioned with members of GH13 subfamilies 17, 23 and 30, of which the prominent activity is α-glucosidase.

**Figure 2.**
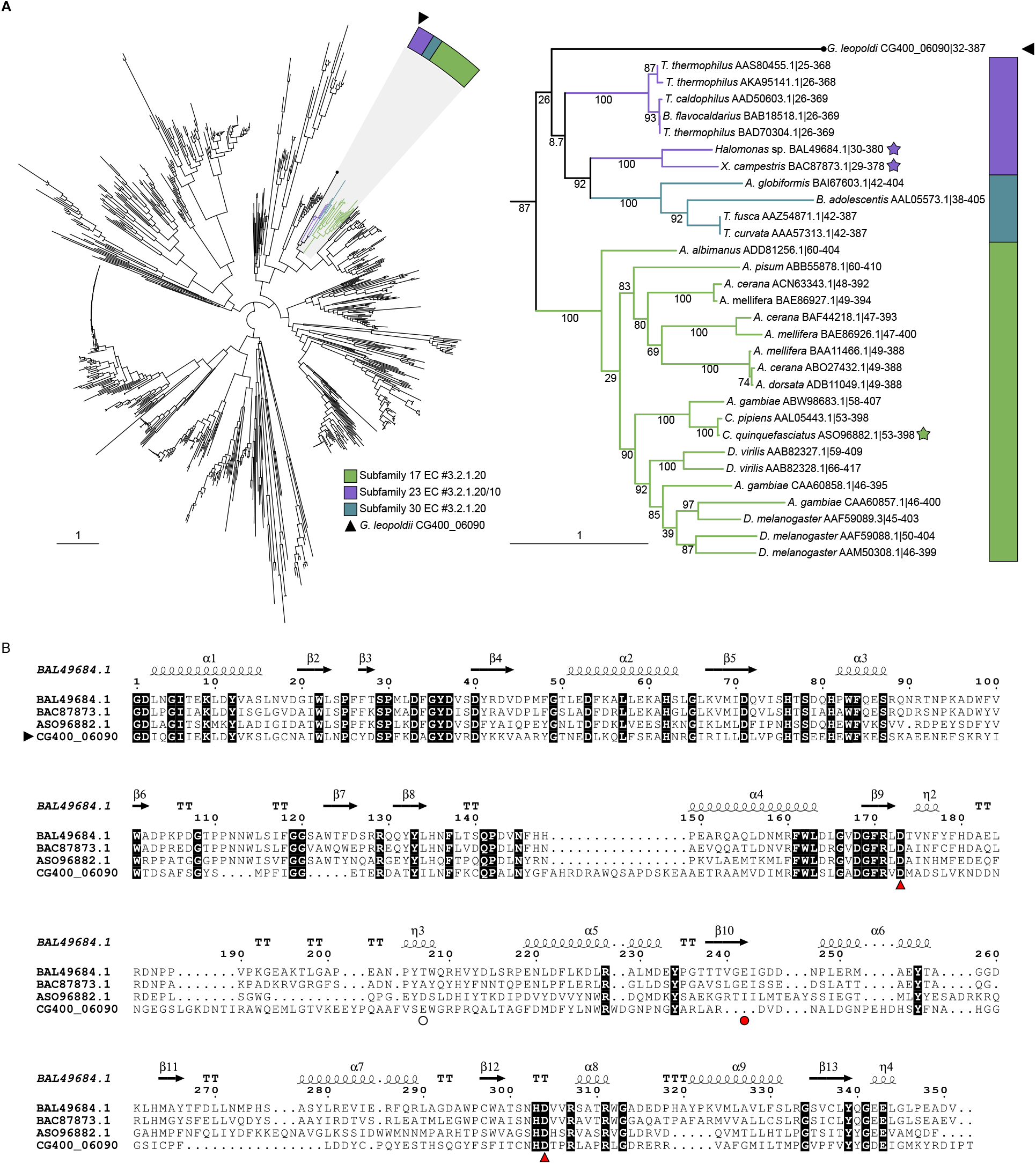
(A) Phylogenetic tree of characterized GH13 members in the CAZy database and predicted *G. leopoldii* NR017 GH13 sequence CG400_06090. (B) Expanded tree of subfamilies 17, 23 and 30. Stars indicate structurally characterized sequences and bootstrap values were included. (C) CLUSTAL alignment of the catalytic domain of CG400_06090 with catalytic domains of GH13 subfamily 23 proteins from *Halomonas* sp. (BAL49684.1) and *Xanthomonas campestris* (BAC87873.1), and subfamily 17 protein from *Culex quinquefasciatus* (ASO96882.1). Red arrows indicate the conserved members of the catalytic triad, and a red circle indicates the acid/base (Glu242) in BAL49684.1 and BAC87873.1. White circle indicates the predicted acid/base in CG400_06090. Invariant sequences are highlighted in black.

This clade was expanded to clarify the relationship of CG400_06090 with related characterized sequences as an apparently deep-branching member of subfamily 23, although with low bootstrap support (Figure 2B). When additional, uncharacterized sequences from the CAZy database were included, CG400_06090 clustered within subfamily 23 (Figure S2).

To identify conserved catalytic residues between CG400_06090 and characterized members of GH13, CG400_06090 was aligned with structurally characterized members of subfamily 23 (BAL49684.1 and BAC87873.1) and subfamily 17 (ASO96882.1) (Figure 2C). Of the GH13 catalytic triad, the aspartate nucleophile and the aspartate that stabilizes the transition state are conserved among the four sequences (39). The glutamate general acid/base, however, does not appear to be sequence conserved between CG400_06090 and the GH13 subfamily 23 members. Upon modelling CG400_06090 in PHYRE2 and aligning with BAL49684.1 (PDBID 3WY1), CG400_06090 Glu256 did appear to be spatially conserved with the general acid/base (not shown).

### Expression and purification of α-glucosidase protein

The *G. leopoldi* NR017 CG400_06090 open reading frame comprises 1701 base pairs encoding a 567 amino acid protein, with a predicted mass of 62 kDa. The full-length open reading frame encoding amino acids 2-567 was PCR-amplified and ligated into vector pQE-80L for expression in *E. coli* as an N-terminal hexahistidine-tagged protein. A distinct protein band in SDS-PAGE between 50 kDa and 75 kDa was obtained from IPTG-induced *E. coli* cells compared with non-induced cells, indicating the expression of the α-glucosidase protein (Figure 3A). The recombinant protein was soluble, and was purified using a Ni-NTA spin column (Figure 3B). From a 50 mL broth culture, approximately 700 μL of purified protein at 4.0 mg/mL was obtained.

**Figure 3.**
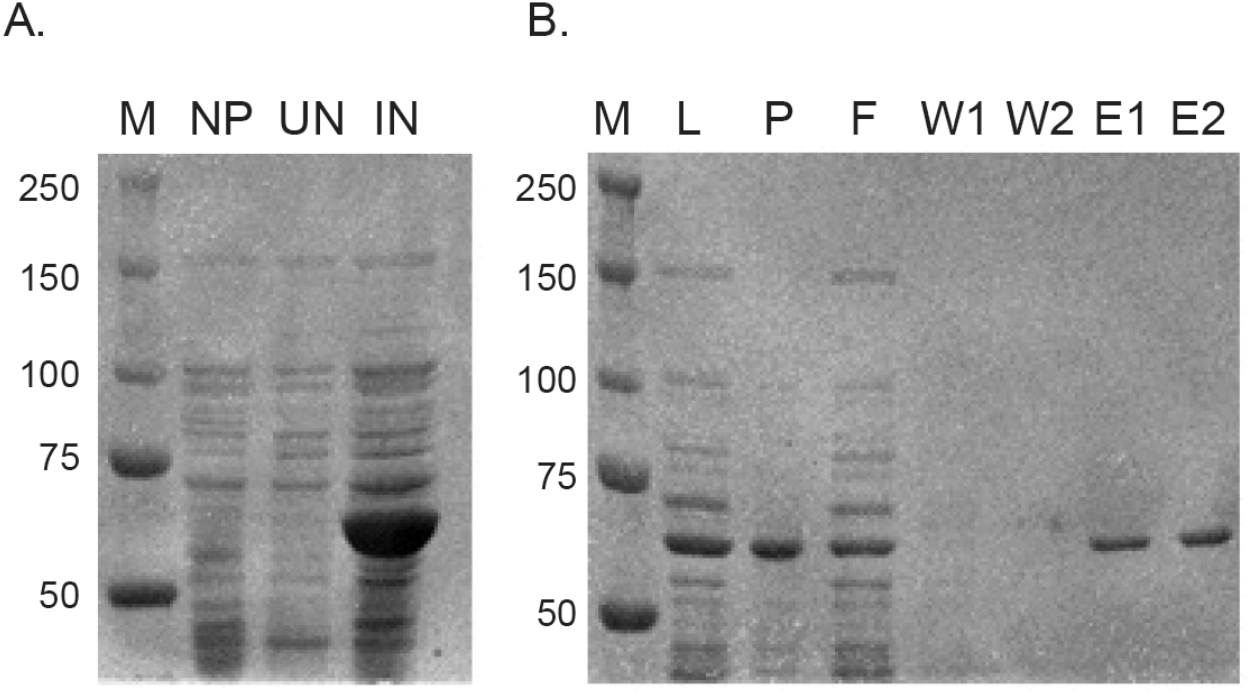
Production and purification of α-glucosidase protein. (A) A protein of the predicted mass of 62 kDa was observed in induced cultures. M: size marker, NP: *E. coli* with no vector, UN: uninduced culture, IN: induced with IPTG. Numbers on the left indicate the marker size in kDa. (B) Fractions from Ni-NTA affinity purification of His-tagged recombinant protein. M: size marker, L: lysate, P: pellet, F: flow-through, W1: first wash, W2: second wash, E1: first elution, E2: second elution.

### Effect of pH on enzyme activity

α-Glucosidase activity of the purified protein was demonstrated by release of 4-nitrophenol from the chromogenic substrate 4-nitrophenyl α-D-glucopyranoside. A preliminary analysis of enzyme activity over pH 3-8 showed that product was produced over a broad pH range from 4.0 to 8.0 (Figure S3). To more precisely examine effects of pH on activity a pH rate profile was determined and the maximum rate was observed at pH 7.0 (Figure 4A). The dependence of rate on substrate concentration fit the Michaelis-Menten equation (Figure 4B), with a Michaelis constant (*K_m_*) of 8.3 μM, and a *V_max_* 96 μM/min, corresponding to a *k*_cat_ value of 0.96 (± 0.01) min^-1^ and *k_cat_/K_m_* of 0.11 μM^-1^min^-1^.

**Figure 4.**
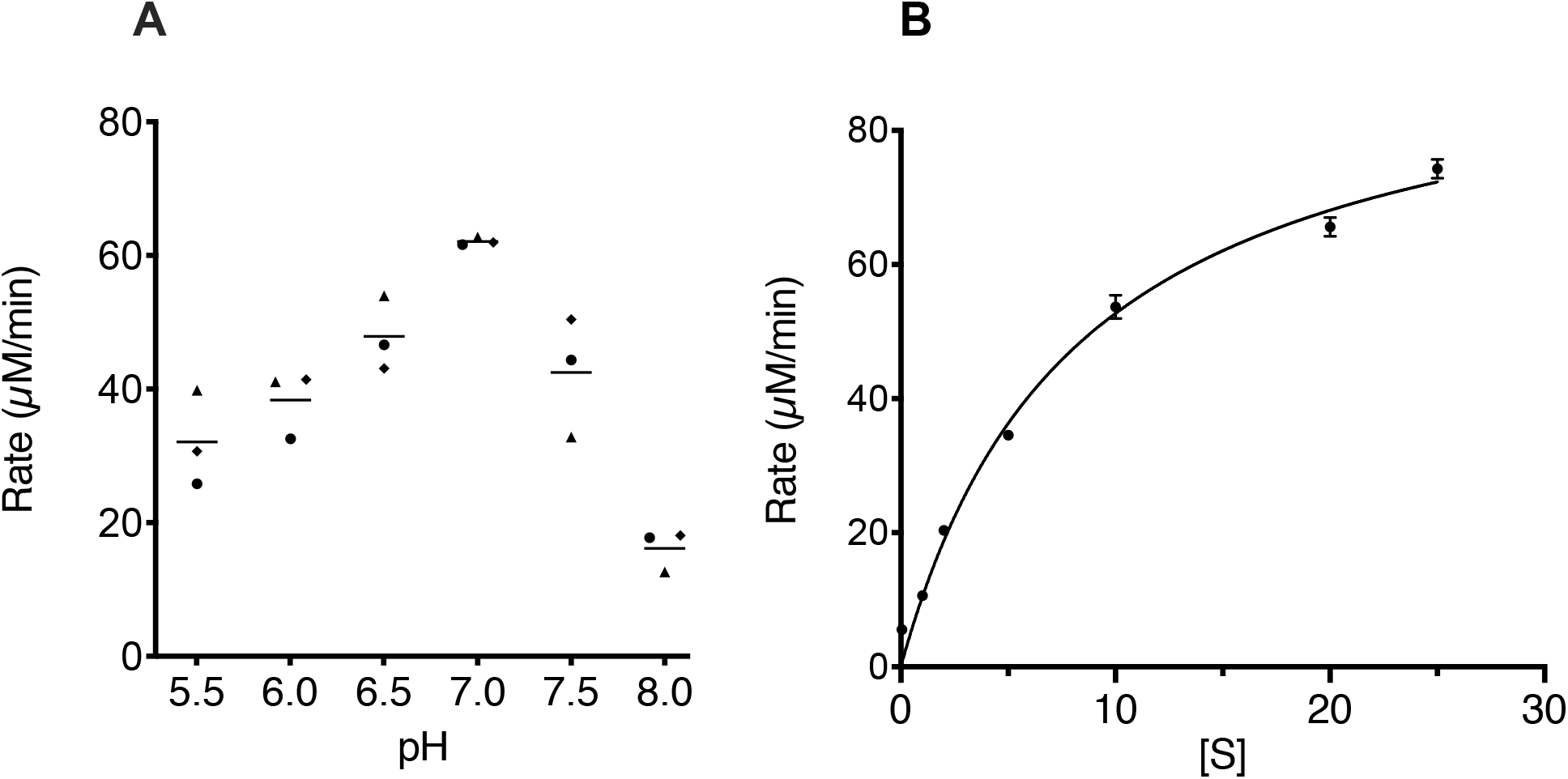
(A) pH-rate profile of α-glucosidase. Each point represents the average of two technical replicates for each of three independent experiments as indicated by different shapes. Horizontal lines indicate the mean. (B) Michaelis-Menten plot. Results shown are the average of three independent experiments, with error bars indicating standard deviation. The line represents the fit to the Michaelis-Menten equation.

### Analysis of substrate hydrolysis

Production of glucose was detected when the purified α-glucosidase was incubated with maltose (M2), maltotriose (M3), maltotetraose (M4) and maltopentaose (M5) (Figure 5). Samples analyzed by HPAEC-PAD contained large peaks corresponding to maltooligosaccharides ranging from maltose to maltopentaose; minor peaks, especially in the maltotetraose and maltopentaose samples, resolved products not observed in the TLC analysis. These peaks did not align to isomaltotriose, panose, or 6-o-a-D-glucosyl-maltose standards in HPAEC-PAD (not shown), and likely represent larger, mixed linkage products. No appreciable activity was detected on maltodextrins (MD 4-7, MD 13-17 and MD 16.5-19.5) or glycogen (Figure S4).

**Figure 5.**
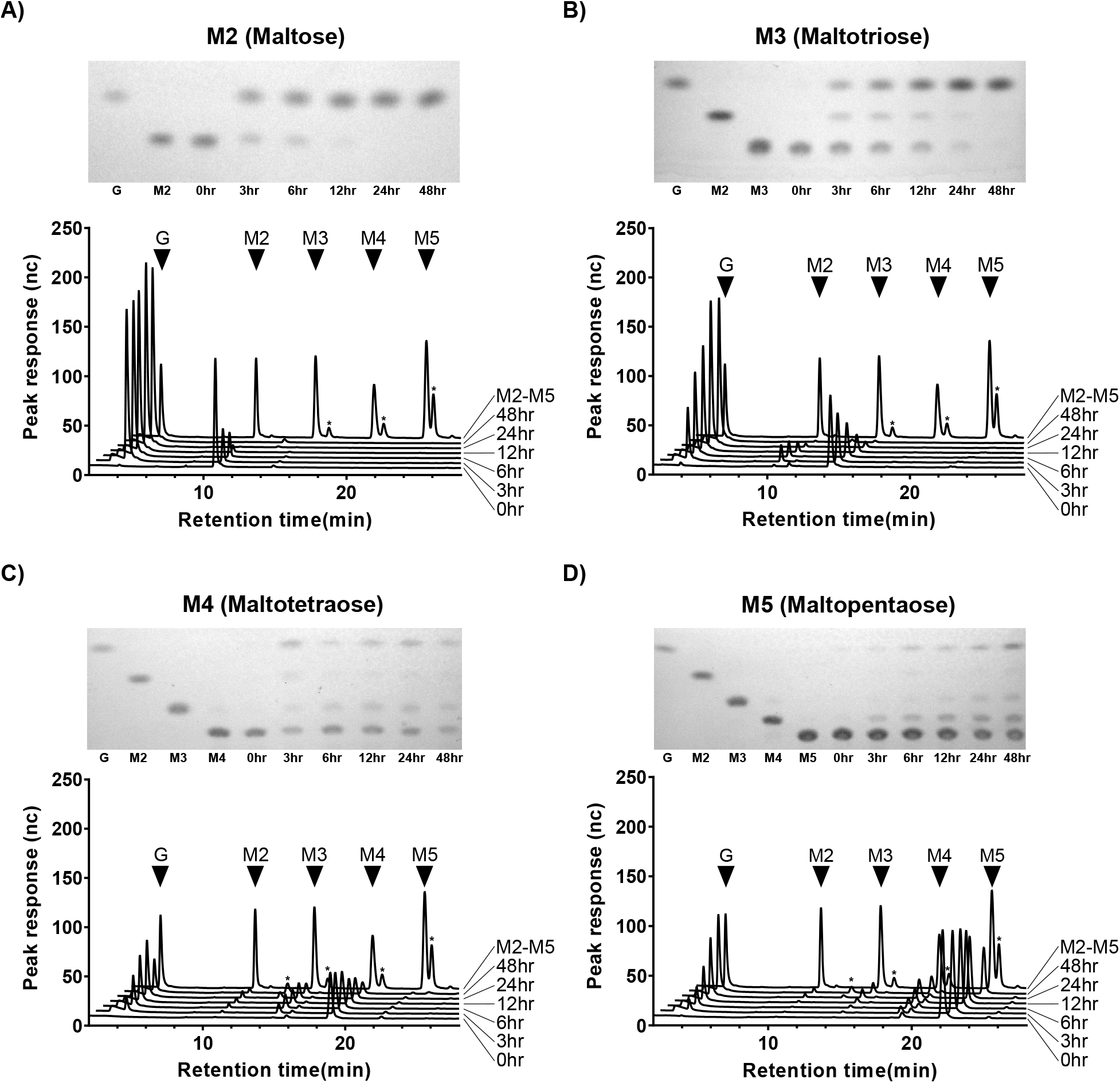
α-glucosidase CG400_06090 digests of maltose to maltopentaose (A-D). Each panel consists of a TLC plate and HPAEC-PAD trace of CG400_06090 digests of M2-M5 between 0hr and 48hr. Major peaks from M2-M5 standards are denoted with black triangles while stars represent unique, undefined oligosaccharides.

## Discussion

Glycogen is a major nutrient available to vaginal microbiota and its utilization likely plays an important role in the survival and success of *Gardnerella* spp. in the vaginal microbiome. The ability of *Gardnerella* spp. to utilize glycogen has been reported in previous studies. Dunkelburg *et al*. described the growth of four strains of what was then known as *Haemophilus vaginalis* in buffered peptone water with 1% glycogen suggesting their ability to ferment glycogen, but no additional details were provided (40). Similarly, Edmunds reported the glycogen fermenting ability of 14 out of 15 *Haemophilus vaginalis* isolates, although serum was included in the media (41). Piot *et al*. examined the starch fermentation ability of 175 *Gardnerella* isolates on Mueller Hinton agar supplemented with horse serum and found that all *Gardnerella* strains were capable of hydrolysing starch (42). They did not, however, assess glycogen degradation and further, it is not clear if serum amylase contained in the media contributed to the observed amylase activity. Robinson, in his review, reported that starch can be utilized by *Gardnerella* (43) while Catlin in another review, reported that glycogen utilization is inconsistent in *Gardnerella* (11).

Glycogen is digested into smaller products, such as maltose and maltodextrins, by a group of enzymes collectively known as amylases. Amylases that contribute to glycogen utilization in the vaginal environment are not well studied. Spear *et al*. (19) suggested a role for a human amylase enzyme in the breakdown of vaginal glycogen. They detected host-derived pancreatic α-amylase and acidic α-glucosidase in genital fluid samples collected from women and showed that genital fluid containing these enzymes can degrade the glycogen. The presence of these enzyme activities alone in vaginal fluid, however, was not correlated with glycogen degradation, suggesting contributions of activity from the resident microbiota. Many vaginal bacteria likely produce amylases that contribute to this activity as various anaerobic taxa are known to encode amylases and digest glycogen (44), however, the contributions of the vaginal microbiota to this process and the importance of these processes to vaginal microbial community dynamics are not yet well understood. Regardless of the source, amylases digest glycogen into maltose and or maltooligosaccharides which are transported inside bacteria via ABC transporter systems for further processing (45).

Protein sequences belonging to eight different GH13 subfamilies that could have roles in glycogen degradation were identified in *G. vaginalis* ATCC 14018 and *G. swidsinskii* GV37 (Figure S1), and multiple amylase-like proteins were detected in the proteomes of additional *Gardnerella* spp. (Figure 1), all of which were annotated as “alpha-amylase” in the GenBank records. Our results demonstrate the value of using more nuanced analyses, including dbCAN and SACCHARIS, to guide discovery of CAZymes based on comparison at a catalytic domain level to functionally characterized enzymes. Further investigation will be required to demonstrate the actual functions of these proteins, and how they contribute to carbohydrate-degrading activities of *Gardnerella* spp..

In the phylogenetic analysis of putative amylase sequences, one cluster contained sequences common to all 26 isolates (Figure 1). Although this protein sequence was annotated as an α-amylase, suggesting it would produce maltooligosaccharides from glycogen, SACCHARIS predicted it to be closely related to α-glucosidases, which cleave glucose from the non-reducing end of its substrate. Automated annotation of sequences deposited in primary sequence databases such as GenBank has resulted in significant problems with functional mis-annotation (46). This problem is exacerbated by the increasing rate at which whole genome sequences are being generated and deposited. The genome sequences of the study isolates were annotated using the NCBI Prokaryotic Genome Annotation Pipeline, which identifies genes and annotates largely based on sequence similarity to previously annotated sequences, thus potentially propagating incorrect functional annotations.

Phylogenetic analysis of GH13 family enzymes showed that the conserved α-glucosidase from *Gardnerella* spp. appears to be most closely related to GH13 subfamily 23 (Figure S2). Primary sequences of GH13 members of amylases have seven highly conserved regions and three amino acids that form the catalytic triad (23). Comparison of the primary sequence and three-dimensional model of CG400_06090 with characterized α-glucosidases from GH13 subfamily 17 and 23 did show sequence conservation of nucleophile and spatial conservation of the catalytic triad.

Our findings suggest that, although α-glucosidase does not hydrolyze glycogen, it can digest smaller maltooligosaccharides. The purified α-glucosidase enzyme was able to completely digest maltose and maltotriose to glucose, but digestion of maltotetraose and maltopentaose was incomplete, suggesting a preference for smaller oligosaccharides (Figure 5). Previously characterized α-glucosidase (*algB*) from *Bifidobacterium adolescentis* DSM20083 also showed higher activity against maltose and maltotriose while no activity was reported against maltotetraose and maltopentaose (47). α-glucosidases can have different substrate specificities due to the variable affinity of the substrate binding site for particular substrates (48).

Most secreted proteins have a short N-terminal signal peptide to guide for extracellular translocation of the newly synthesized protein (49), however, bacterial proteins can be also secreted via non-classical secretion pathways in a Sec-independent manner, without having a signal peptide (28). Analysis of the *Gardnerella* α-glucosidase sequence with SignalP and SecretomeP showed that the protein is unlikely to be secreted by either route and thus likely acts on intracellular substrates including products of extracellular glycogen degradation transported into the cell. Many free-living and host-associated bacteria synthesize glycogen as a storage molecule (50) but the extent to which this occurs in *Gardnerella* is not known. The possibility that products of digestion of intracellular glycogen stores are substrates for the α-glucosidase characterized in this study remains a question for future study. Purified α-glucosidase from *G. leopoldii* NR017 was active across a broad pH range of 4.0 to 8.0 (Figure S3) with the highest rate detected at pH 7.0 (Figure 4A). This is not unexpected for a cytoplasmic enzyme in a host-associated bacterium, and consistent with at least one other report of an α-glucosidase from a commensal bacterium, *Bifidobacterium adolescentis* (47).

Our findings show that *Gardnerella* spp. have an α-glucosidase enzyme that likely contributes to the complex and multistep process of glycogen utilization by releasing glucose from maltooligosaccharides. Identification and biochemical characterization of additional enzymes involved in glycogen metabolism will provide insight into whether utilization of this abundant carbon source is an important factor in population dynamics and competition among *Gardnerella* spp.. The functional annotation strategy demonstrated here provides a powerful approach to guide future experiments aimed at determining enzyme substrates and activities.

## Acknowledgements

The authors are grateful to Josseline Ramos-Figueroa, Douglas Fansher and Natasha Vetter (Department of Chemistry, University of Saskatchewan) for guidance with thin layer chromatography and Champika Fernando for technical support.

---

**Figure S1.**
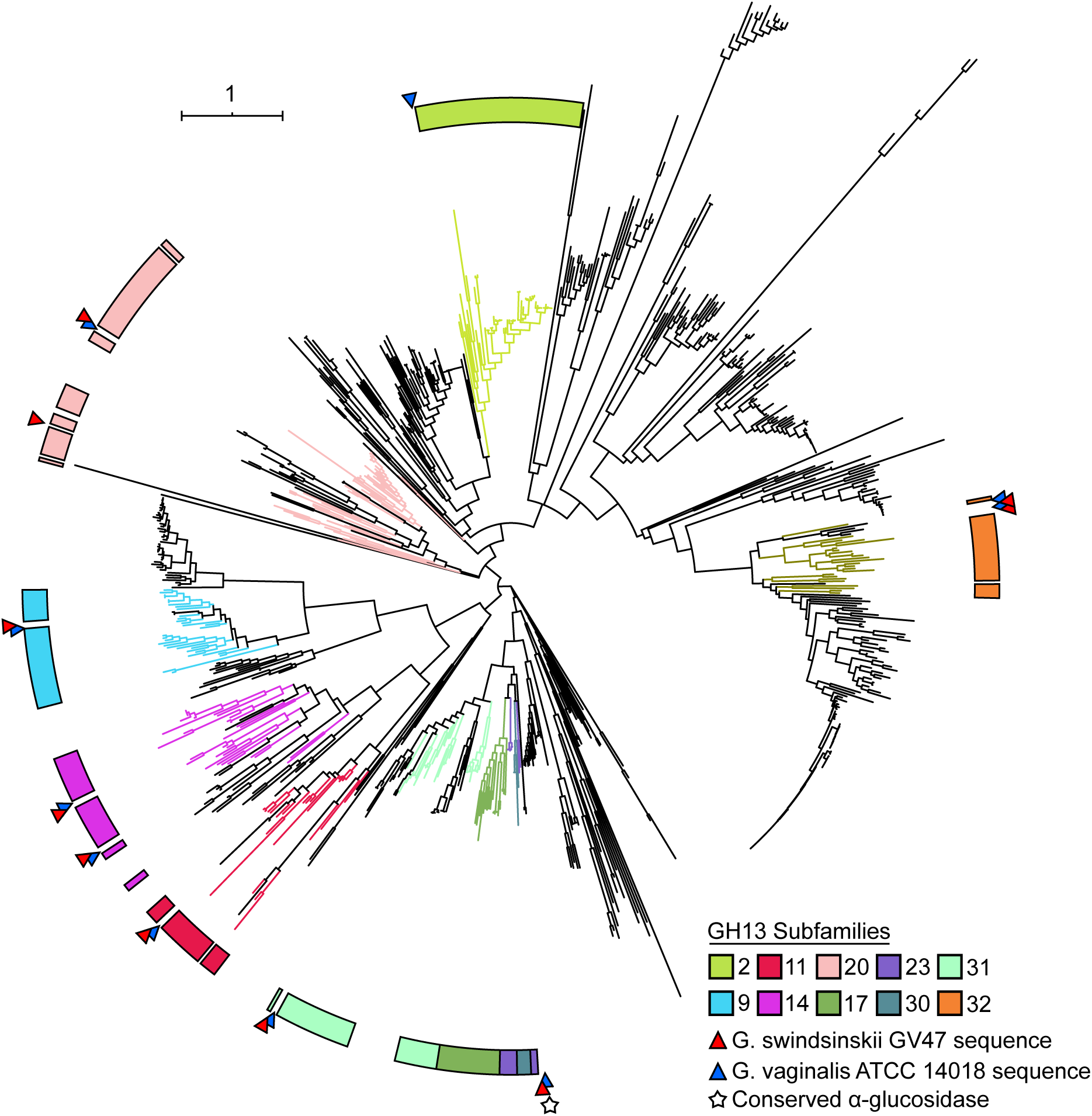
Phylogenetic tree of GH13 functionally characterized members with predicted GH13 domains from *G. swindsinskii* GV37 (red arrow) and *G. vaginalis* ATCC 14018 (blue arrow). Tree branches are coloured based on GH13 subfamily. The conserved α-glucosidase between the 26 proteomes is denoted by a white star. Trees were generated using SACCHARIS and viewed in iTOL.

**Figure S2.**
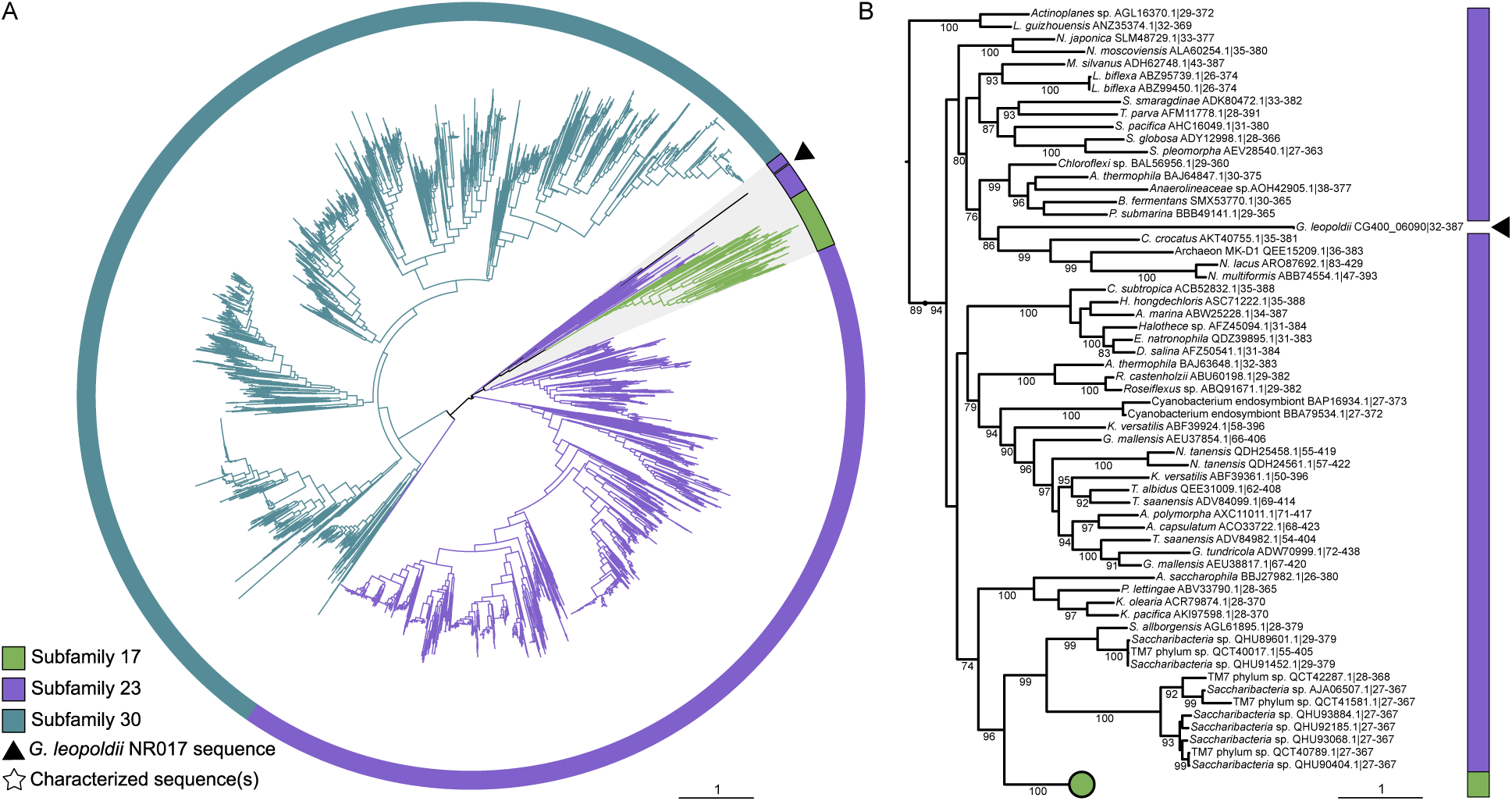
(A) Phylogenetic tree of all GH13 subfamily 17, 23 and 30 members in the CAZy database and *G. leopoldii* CG400_06090. Branch colour is based on GH13 subfamily and functionally characterized proteins are indicated with stars. The clade highlighted in grey including *G. leopoldii* CG400_06090 and its closest relatives is expanded in (B).

**Figure S3.**
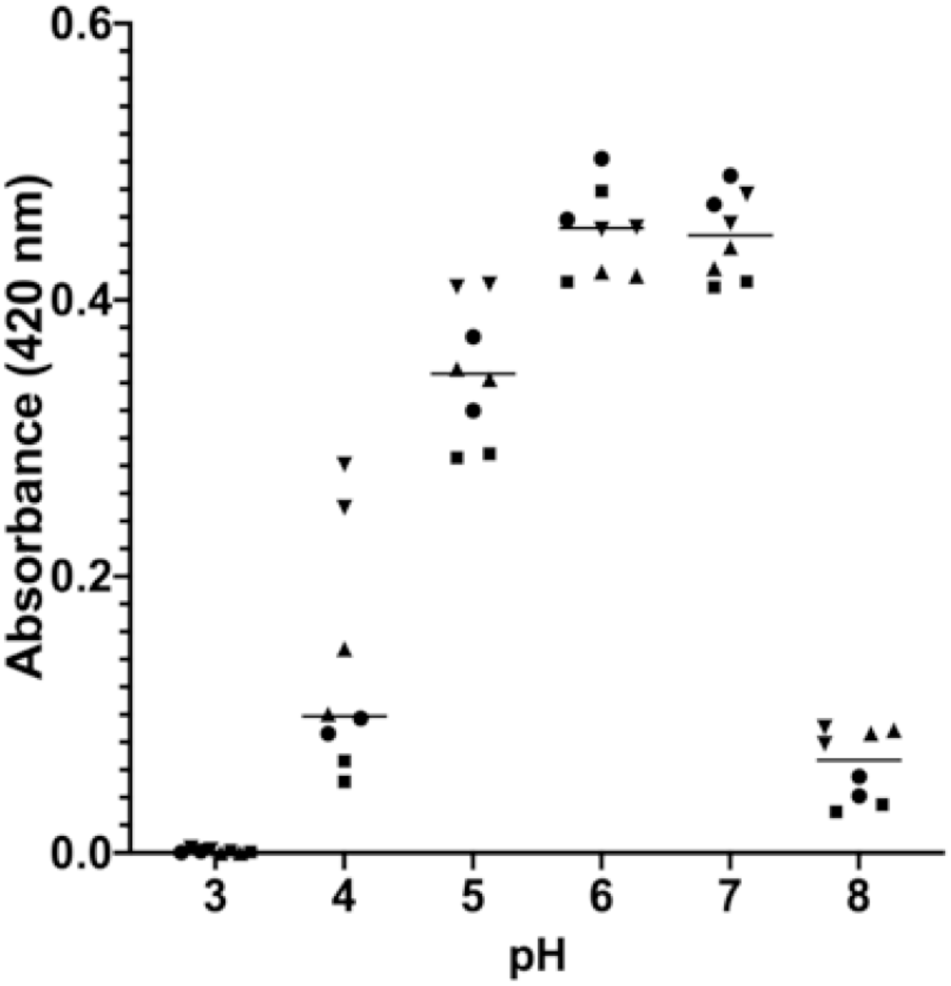
Release of 4-nitrophenol from chromogenic substrate 4-nitrophenyl-α-D-glucopyranoside at pH 3-8. Results from four independent experiments each with two technical replicates are shown. The α-glucosidase (0.8 mM) was incubated with 10 mM substrate at different pH and amount of 4-nitrophenol released in 10 minutes was measured.

**Figure S4.**
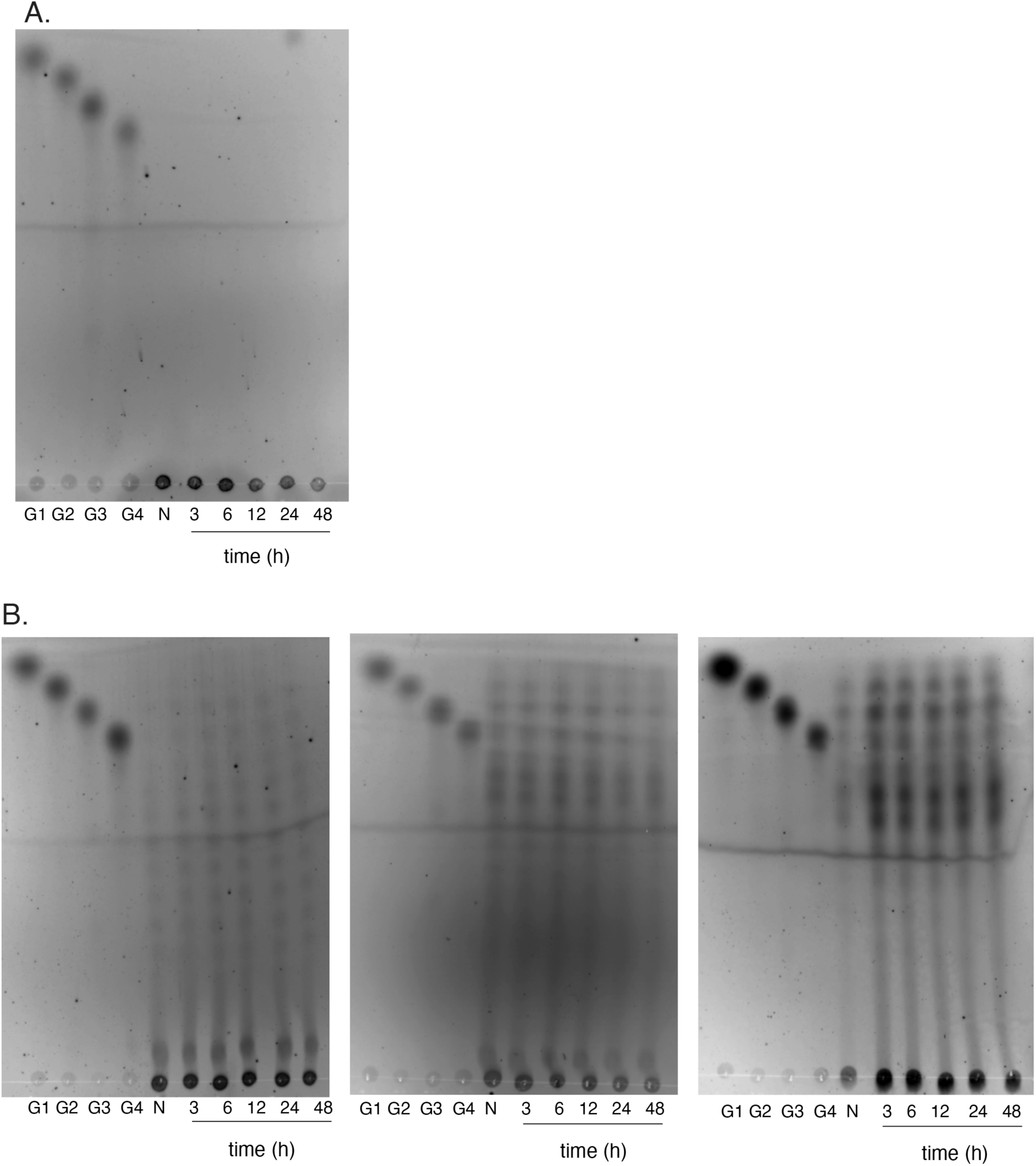
(A) TLC of products of glycogen hydrolysis by α-glucosidase enzyme. Reaction mixtures were assessed at 3 h, 6 h, 12 h, 24 h and 48 h. N = Substrate with no enzyme. (B) TLC of products hydrolysis by α-glucosidase enzyme of maltodextrins MD 4-7 (left panel), MD 13-17 (middle panel) and MD 16.5-19.5 (right panel). Reaction mixtures were assessed at 3 h, 6 h, 12 h, 24 h and 48 h. N = Substrate with no enzyme.

